# 3D Genome Structure Variation Across Cell Types Captured by Integrating Multi-omics

**DOI:** 10.1101/784223

**Authors:** Yang Xu, Tongye Shen, Rachel Patton McCord

## Abstract

**Background:** 3D genome structure contributes to the establishment or maintenance of cell identity in part by organizing genes into spatial active or inactive compartments. Less is known about how compartment switching occurs across different cell types. Rather than analyze individual A/B compartment switches between pairs of cell types, here, we seek to identify coordinated changes in groups of compartment-scale interactions across a spectrum of cell types.

**Results:** To characterize the impact of genome folding on cell identity, we integrated 35 Hi-C datasets with 125 DNase-seq, 244 RNA-seq, and 893 ChIP-seq datasets. We first find physical associations with the nuclear lamina inform the most dramatic changes in chromosome structure across cell types. By examining variations in chromosome structure, transcription, and chromatin accessibility, we further observe that certain sets of correlated chromosome structure contacts also co-vary in transcription and chromatin accessibility. Analyzing ChIP-seq signals, we find that sets of chromosome contacts that form and break in sync tend to share active or suppressive histone marks. Finally, we observe that similar principles appear to govern chromosome structure fluctuations across single cells as were found across cell types.

**Conclusion:** Our results suggest that cells adapt their chromosome structures, guided by variable associations with the lamina and histone marks, to allocate up-regulatory or down-regulatory resources to certain regions and achieve transcription and chromatin accessibility variation. Our study shows E-PCA can identify the major variable interaction sets within populations of single cells, across broad categories of normal cell types, and between cancer and non-cancerous cell types.

## Background

It is commonly accepted that human cell types can be classified by their different patterns of gene expression and epigenetic status. Indeed, numerous recent studies use this idea to classify different cell types in tissues like brain and kidney, for example, by single cell RNA sequencing, chromatin accessibility and state profiling [1], [2], [3]. In addition to gene expression and epigenetic state, the 3D folding of the genome also appears to play a crucial role in defining cell type. From the first Hi-C dataset, it was clear that a major variation in 3D genome structure between cell types is the pattern of chromosome spatial compartmentalization into “A” (active) and “B” (inactive) compartments [4]. Indeed, in a more recent compendium of Hi-C maps across human cell types, it was observed that about 60% of genomic loci are “dynamically” compartmentalized in different tissues and cell types [5]. When regions change compartments, this shift is often associated with a change in expression of genes in those regions [6]. As a consequence, numerous studies have tried to characterize the differences in spatial compartmentalization between cell types and its relationship to other linear features of the genome [7], [8], [9], [10]. So far, however, the most common analyses track changes between pairs of cell types or conditions in terms of which regions switch their compartment assignment from A to B or where the boundaries of topologically associating domains (TAD) are strengthened or lost.

With the proliferation of Hi-C datasets, approaches are needed to simultaneously compare the patterns of genome structure change across many datasets at once and to identify the major patterns of change. Methods are also needed to move beyond identifying isolated changes of compartmentalization in certain loci and instead to identify how chromosome contacts change coordinately across the chromosome. Several studies have tried to elucidate cell-to-cell variation within chromosome structure. For instance, Sauerwald and Kingsford used the sample-pair comparison approach to identify similar TAD intervals across 23 human Hi-C samples and found cancer cells show higher structural variability in pan-cancer gene regions [11]. However, two-sample comparison methods may not recognize an emergent collective pattern that arises from a comprehensive comparison of multiple cells, and pairwise comparisons do not reveal whether detected differences are statistically correlated. Some studies also have used PCA to compare A/B compartment profiles across cell types [10], but such analyses still require first converting 2D Hi-C profiles into 1D profiles, and then using the 1D information to infer cellular variations. Such analysis does not analyze what sets of variations drive the segregation of different cell types.

We previously reported the application of a contact matrix analysis originally developed for the study of a protein’s structural dynamics, E-PCA, to a set of Hi-C matrices across nine different blood cell types on a portion of one chromosome [12]. Our results showed that the strength of compartment segregation was the major source of variation across those cell types, but we did not further investigate which particular A-A or B-B associations formed or separated coordinately. Here, we apply E-PCA to 39 Hi-C datasets across widely varying cell types to determine, across all chromosomes, which interactions contribute most to the chromosome structure fluctuation across cell types. We identify regions that exhibit coordinated structure change: that is, certain regions switch their interaction partners in concert across cell types. When one set of interactions breaks, another set forms. We find that lamina associated domains (LADs) are a major genomic feature that correlates with the coordinated changes in compartmentalization among cell types. Further, there appears to be a coordinated switch between long distance associations mediated by the lamina and interactions in a shorter range involving super-enhancers. The interactions that define the major modes fluctuation tend to occur at genomic regions that are highly variable in their transcription and chromatin accessibility. We further find that patterns of chromosome structure fluctuation are related to certain sets of histone modifications. Strikingly, these underlying factors that appear to guide the major fluctuations across cell types also are found to be associated with the major fluctuations in chromosome contacts across individual cells in single cell Hi-C datasets.

## Results

### Analysis pipeline for the elucidation of genome structure variation

We present our main analysis pipeline in Fig. 1. We first collected Hi-C datasets that cover a variety of cell types which are available from previous publications, ENCODE, or our own laboratory, and then we processed them into interaction contact matrices at 100kb resolution. We next applied E-PCA to locate key sets of interactions that vary coordinately across cell types (step 2). The details of the E-PCA approach can be found in our previous study [12]. Briefly, E-PCA takes as an input a set of contact matrices (here, Hi-C data from different cell types), flattens each matrix into a vector of contacts, combines all these contact vectors into one matrix that contains every pairwise chromosome contact (rows) across all input cell types (columns), and then uses PCA to detect correlated patterns of contacts (PC1, PC2, etc.) that best capture the variation across cell types. We isolated the bin pairs in the top 1% of positive and separately top 1% of negative PC loadings as “key variable interactions” and classified these genomic regions according to their nuclear lamina association annotation (step 3). We also represent this top 1% of key variable interactions in circular plots (step 4). This circular plot intuitively shows the interaction switches that occur between positive and negative PC scenarios. It also allows clear annotation of nuclear lamina association, transcription variation, or chromatin accessibility variation that characterize these positive and negative interaction sets. In step 5, we filtered the key interactions from step 4which link pairs of regions that are co-transcribed and/or co-accessible according to RNA- and DNase-seq data across cell types. This allows us to identify which sets of interactions are most related to cell-type specific gene expression changes or local chromatin state changes. In step 6, we scored each genomic bin by how often it is involved in a key variable interaction and use this 1D information to compare with peaks called from ChIP-seq that targets different histone marks. Our pipeline takes advantage of E-PCA to extract chromosome interaction sets that can be integrated with other 1D regulatory signals. We used the approach to elucidate the variation of genome structure and successfully interpreted the observed variation from different angles.

**Fig. 1.**
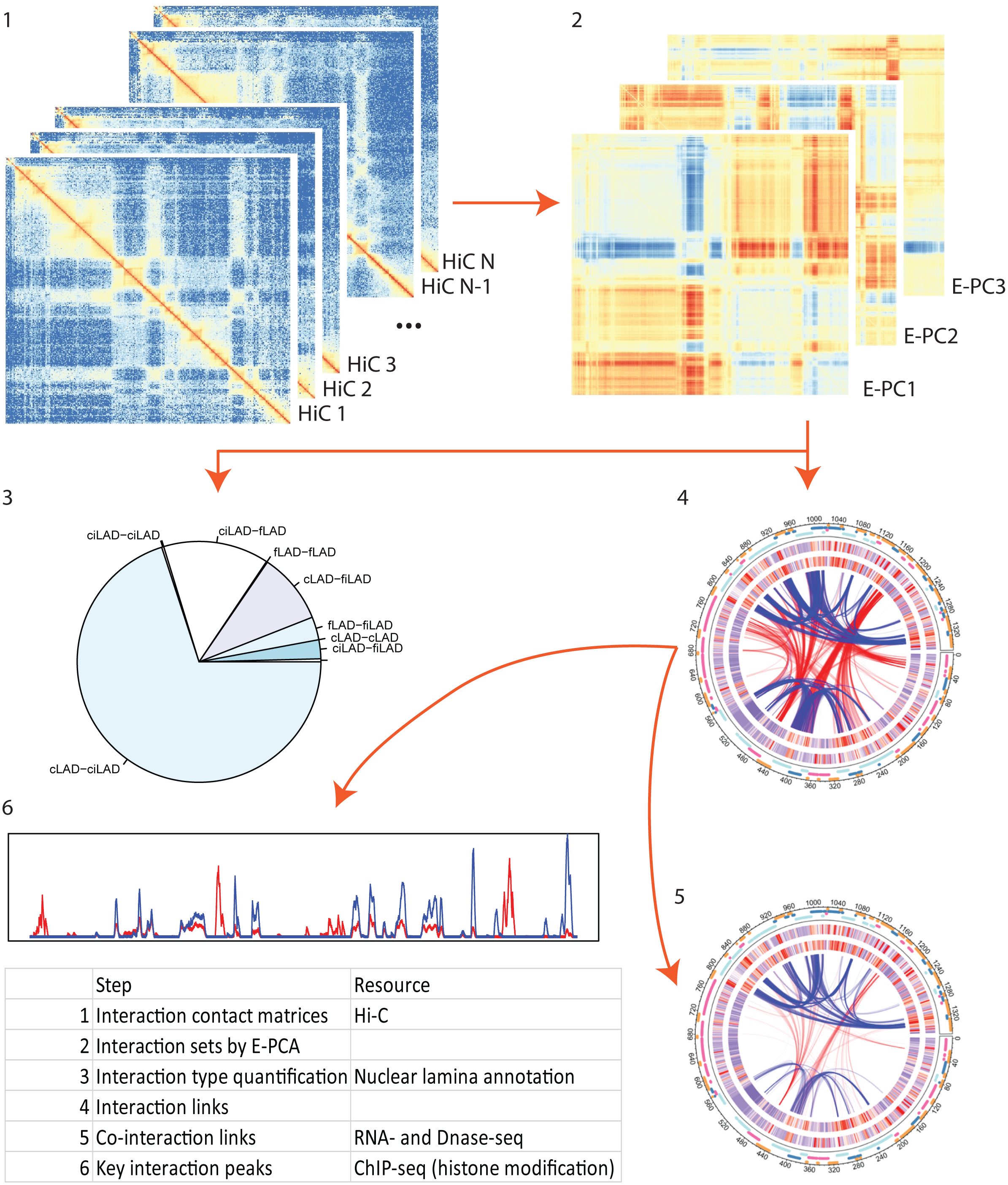
A schematic of the analysis workflow. See full description of workflow in main text.

### E-PCA captures coherent chromosome structural variance

Principal component analysis (PCA) on Hi-C chromosome contact matrices can reveal consensus features and major structural variations. Specially, the conventional chromosome contact PCA highlights the spatial separation of active and inactive chromatin regions but is unable to dissect the major fluctuations of chromosome structure across different cell types. Our previous work applying E-PCA to microscopy data and Hi-C data demonstrated an alternative approach to capture dominant motions of chromosome structure [12]. Here, we apply this E-PCA approach to find major variations of genome structure for each chromosome across 35 Hi-C datasets from different cell types (Table 1). Among these 35 datasets, we included two to three replicates for some cell types. This allows us to verify that the important variations reported here are not from technical differences. We also include both primary cells and cell lines, cancer, blood, fibroblast, and epithelial types. While this is not a comprehensive set of cell types, our focus is to demonstrate how E-PCA can derive general principles of chromosome structure changes across a diverse range of samples.

**Table 1.** Summary of Hi-C samples. Samples are indexed from 1 to 35 with brief description and data source.

| Index | Cell type abbreviation | General type | Culture type | Cell type description | Source |
| --- | --- | --- | --- | --- | --- |
| 1 | Hepatic endothelial | Endothelial | Primary cell | Primary endothelial cell of hepatic sinusoid | ENCSR982KWR |
| 2 | DLD1 | Cancer | Cell line | colorectal adenocarcinoma | ENCSR213DHH |
| 3 | A375 | Cancer | Cell line | skin malignant melanoma | In-house |
| 4 | A549_r1 | Cancer | Cell line | lung adenocarcinoma, epithelial | ENCSR444WCZ |
| 5 | A549_r2 | Cancer | Cell line | lung adenocarcinoma, epithelial | ENCSR444WCZ |
| 6 | HepG2_r1 | Cancer | Cell line | hepatocellular carcinoma | ENCSR194SRI |
| 7 | HepG2_r2 | Cancer | Cell line | hepatocellular carcinoma | ENCSR194SRI |
| 8 | SJCRH30 | Metastatic Cancer | Cell line | metastasized to bone marrow | ENCSR998ZSP |
| 9 | SKNDZ | Metastatic Cancer | Cell line | neuroblastoma, metastasized to bone marrow | ENCSR105KFX |
| 10 | SKNMC | Metastatic Cancer | Cell line | neuroepithelioma, metastasized to supra-orbital | ENCSR834DXR |
| 11 | T47D | Metastatic Cancer | Cell line | ductal carcinoma, metastasized to lung | ENCSR549MGQ |
| 12 | Erythroblast | Blood | Primary cell | Erythroblast | Javierre et al. |
| 13 | Macrophage | Blood | Primary cell | Macrophage | Javierre et al. |
| 14 | MK | Blood | Primary cell | MK | Javierre et al. |
| 15 | Monocyte | Blood | Primary cell | Monocyte | Javierre et al. |
| 16 | B cell | Blood | Primary cell | B cell | Javierre et al. |
| 17 | CD4+ cell | Blood | Primary cell | CD4+ cell | Javierre et al. |
| 18 | CD8+ cell | Blood | Primary cell | CD8+ cell | Javierre et al. |
| 19 | Neutrophil | Blood | Primary cell | Neutrophil | Javierre et al. |
| 20 | Hffc6_r1 | Fibroblast | Cell line | foreskin fibroblast | In-house |
| 21 | Hffc6_r2 | Fibroblast | Cell line | foreskin fibroblast | In-house |
| 22 | Cerebellum astrocyte | Astrocyte | Primary cell | Astrocyte of the cerebellum | ENCSR011GNI |
| 23 | Brain endothelial | Endothelial | Primary cell | Brain microvascular endothelial cell | ENCSR507AHE |
| 24 | Brain pericyte | Pericyte | Primary cell | Brain pericyte | ENCSR323QIP |
| 25 | HGADFN167 | Fibroblast | Primary cell | Progeria patient fibroblast | In-house |
| 26 | Endometrial endothelial | Endothelial | Primary cell | Endometrial microvascular endothelial cell | ENCSR551IPY |
| 27 | ACHN | Metastatic Cancer | Cell line | renal cell adenocarcinoma, metastasized to lung | ENCSR343VKT |
| 28 | Spinal Cord Astrocyte | Astrocyte | Primary cell | Astrocyte of the spinal cord | ENCSR974LXZ |
| 29 | ATMhTERT | Fibroblast | Cell line | ATM mutant fibroblast, hTERT immortalized | GSE136899 |
| 30 | BJ1 | Fibroblast | Cell line | foreskin fibroblast, hTERT immortalized | GSE136899 |
| 31 | GM02052 | Fibroblast | Cell line | ATM mutant fibroblast | GSE136899 |
| 32 | GM12878_r1 | Blood | Cell line | GM12878_r1 | ENCSR968KAY |
| 33 | GM12878_r2 | Blood | Cell line | GM12878_r2 | GSE136899 |
| 34 | GM12878_r3 | Blood | Cell line | GM12878_r3 | GSE136899 |
| 35 | MRC5 | Fibroblast | Cell line | fetal lung fibroblast | GSE136899 |

E-PCA reflects not only which interactions vary across cell types, but how those varying interactions are correlated, and thus may work together to drive variation among cell types. The E-PCA results presented in Fig. 2a illustrate that positive interactions (red) usually form as a set in the same cell type with other positive interactions (red), and that these interactions tend to break when the set of negative interactions (blue) forms. We focus on the first three principal components labeled E-PC1 – E-PC3, as these capture at least 70% of the variation across Hi-C datasets in most chromosomes (Table S1). We focus on a few chromosomes as examples in the main figures, but chromosome structural variations captured by E-PCA for all chromosomes are shown in Figure S1. The first three principal components capture distinct types of structure variation and suggest a high complexity in chromosome folding (Fig. 2a). It is important to note that the overall sign of positive and negative PC values (indicated in red and blue) are arbitrary, only showing interactions that are anti-correlated (when red interactions form, blue interactions break and vice versa). As such, the types of interactions labeled positive and negative by E-PCA for one chromosome may have an inverted sign on another chromosome.

**Fig. 2.**
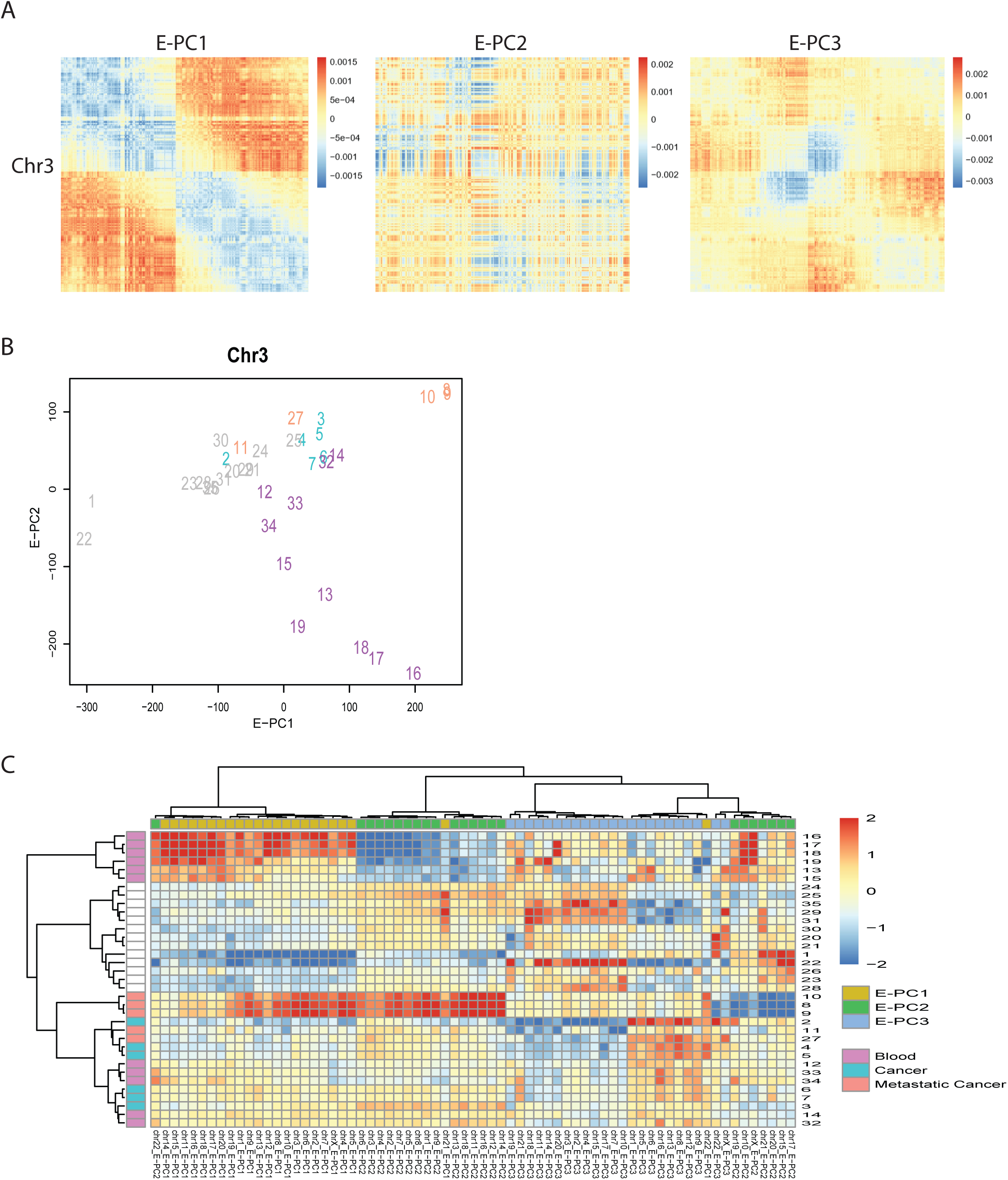
Chromosome folding variation captured by E-PCA. a. Three major modes of structure variation from chr3 are represented by the first three E-PC loadings. b. Scatter plot of 36 Hi-C datasets projected onto the first two PCs. c. Hierarchical clustering of PCs (columns) and datasets (rows) performed across the first 3 E-PC projections across all 23 chromosomes from the 36 samples. The separation of different cell types is largely consistent across different chromosomes.

With E-PCs representing the major modes of chromosome structure fluctuation across the 35 Hi-C samples, we can project each sample into the first two principal component spaces to identify which cell types are most similar to and different from one another with respect to these modes of variation (Fig. 2b and Figure S2). For example, the chromosome structure variations captured by chr3 E-PC2 separate blood cell types from fibroblasts, epithelial, and endothelial cells. We next investigated whether the top E-PC patterns generated from one chromosome have any relation to those from another chromosome. To generate a consensus view of cell type grouping, we performed hierarchical clustering on the projections of each cell type onto the first three E-PCs across all chromosomes. Strikingly, we found most E-PC1s were clustered into one group, with separate clusters emerging for E-PC2s and E-PC3s (Fig. 2c). PCA on each chromosome was performed independently and thus there is no mathematical reason that E-PC1 from different chromosomes would necessarily segregate the cell types in a similar way. The fact that E-PC1, 2, and 3 projections form distinct clusters indicates chromosome structural variations displayed by each principal component are capturing a similar biological mode across chromosomes. However, there are some exceptions. For instance, chr21 E-PC3 was clustered with E-PC1s of other chromosomes, suggesting variations captured by chr21 E-PC3 is like variations in E-PC1s of other chromosomes. Examining the clustering patterns of the cell types, we found that 6 blood cell types, including B cell, CD4+ T cell, CD8+ T cell, Neutrophil, Macrophage and Monocyte, are clustered in a group, fibroblasts (two Hff replicates and BJ1) cluster together, and two A549 lung cancer cell replicates are closely clustered with other cancer cell lines (T47D, ACHN, A375, and two HepG2 replicates). The results above indicate two perspectives for us to understand chromosome structure variation between cell types: structure modes of different chromosomes follow similar principles and 2) similarities or differences between cell types can be distinguished by interaction sets rather than simply by individual interactions or compartment switches.

### Nuclear lamina interaction associates with the most prominent mode of chromosome structure change across cell types

After defining the correlated patterns of interaction variation across cell types, we next sought to identify biological features along the linear genome that could help explain these patterns. Genomic regions often have contacts with the nuclear lamina (lamina associated domains, termed LADs). LADs help anchor interphase chromosome topology and play a role in gene regulation [13], [14]. Regions commonly associated or rarely associated with the lamina have different levels of gene expression and distinct histone marks [15]. A modified DamID method has been used to characterize genome-nuclear lamina interactions across cell types at a single cell level, classifying genomic regions according to whether they associate with lamina always, never, or sometimes [16], [17]. We used this genome-wide LAD annotation to investigate whether chromosome structure variations captured by E-PCA are associated with physical attachment with nuclear lamina.

LADs have been classified by Kind *et al*. as “constitutive” (cLAD), genomic regions associated with the lamina in nearly every cell type, and “facultative” (fLAD), genomic regions that change their lamina association in different cell types. We assigned the LAD type to each 100kb genomic bin and compared these annotations with key variable interactions in heatmaps and circular plots (Fig. 3) [17]. Chromosome 21 provides a clear example of the result we found consistently across other chromosomes. The chromosome contact fluctuations captured by E-PC1 of chr21 involve changes in interactions between the genomic regions annotated as fLAD (facultatively associated with the nuclear lamina) and the rest of the chromosome. One set of highly correlated interactions, captured by positive E-PC1 values, are revealed to be coordinated interactions of fLAD regions with ciLAD regions (genomic bins that are constitutively located in the nuclear interior). Interactions with strong negative E-PC1 values occur between the same fLADs and cLAD regions. This indicates that the major structural switch across the 35 Hi-C datasets on chromosome 21 involves fLADs interacting mutually exclusively either with LAD-associated or interior regions on the rest of the chromosome. (Fig. 3a and 3b). To capture this type of switch quantitatively across all chromosomes, we classified all E-PC positive and negative interactions according to their LAD interaction type. We found that this type of LAD interaction switch is the major fluctuation across most chromosomes. As stated above, for chr21, fLAD-ciLAD interactions represent most positive E-PC1 interactions, while fLAD-cLAD characterize negative E-PC1 interactions (Fig. 3c). For the remaining 22 chromosomes, we also found that one or two LAD interaction types account for the major interactions in E-PC1s and E-PC2s. Specifically, many E-PC2s are a tradeoff between cLAD-ciLAD and cLAD-c/fLAD. While E-PC2 is more related to folding of cLADs, E-PC1 is more related to opening/closing of fLAD regions. (Fig. 3b and Figure S3). In summary, we reason that the major source of cell-to-cell variation in chromosome structure is attachment or detachment of a set of regions from the nuclear lamina.

**Fig. 3.**
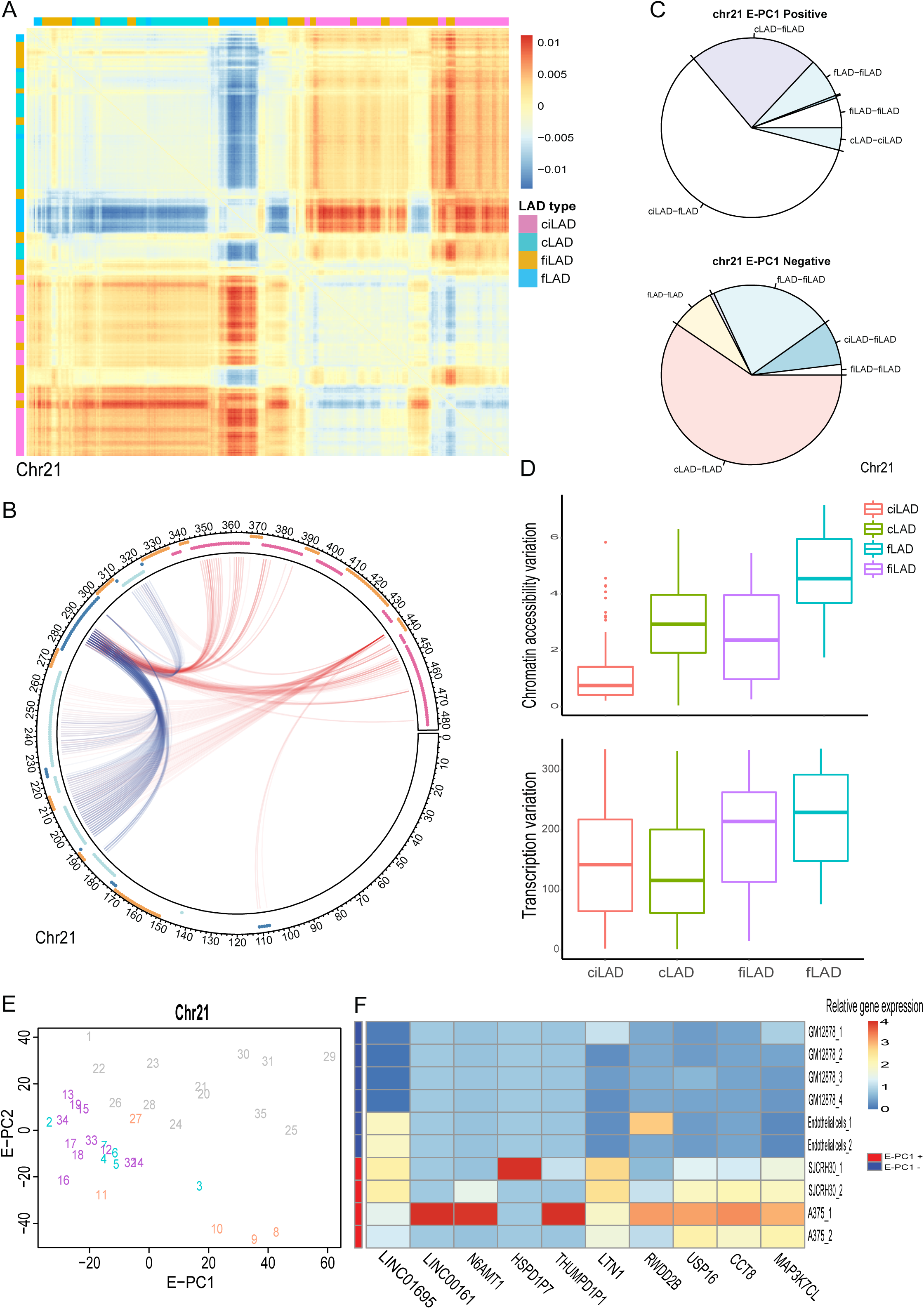
Nuclear lamina association is the major source of variation for chromosome folding. a. Heatmap shows positive and negative interaction sets identified in chr21 E-PC1. The nuclear lamina association type of each genomic bin was added as a track aside the heatmap. b. Pie charts quantify the proportion of each lamina interaction type for positive and negative interaction sets, respectively. c. The circular plot presents top 10% key variable interactions from each positive and negative set of E-PC1. Each link bridges two genomic bins, and the annotation of lamina type is added as a track outside the circular plot. Coordinates indicate the number of 100 kb bins. d. The boxplot quantifies chromatin accessibility and expression variation across chr21 for each lamina association type. e. Scatter plot of 39 Hi-C datasets projected onto the first two PCs of chr21. f. Scaled relative gene expression of genes in the chr21 region that experiences the most variable interactions. Expression level was scaled from 0 to 4 and 0 represented the lowest expression level among the 10 quantified genes.

### Lamin-associated chromosome structure fluctuations coincide with cell type specific gene expression

Chromosome structure changes across cell types may influence cell-type specific gene regulation. Indeed, the relationship we observed between major chromosome structure fluctuations and lamin association suggests a connection between structure variation and gene expression, since lamin association is related to gene repression. For example, we would hypothesize that the major interaction switch observed for E-PC1 on chr21 results in activation of the fLAD region when it is in contact with the ciLADs (red links) and repression of the fLAD region when it is in contact with cLADs (blue links) (Fig. 3b) [15]. We explicitly tested this predicted implication of chromosome structure for gene expression by identifying the 4 cell types with the most extreme E-PC1 projections which also have available RNA-Seq data (Fig. 3e). We examined gene expression in the fLAD regions that exhibited the large interaction switch among cell types: from chr21:28,100,000 to chr21:29,000,000. fLADs in this region interact with repressive cLADs in GM12878 and endothelial cells, while the same fLADs switch to interacting with ciLADs in SJCRH30 and A375. This lamina association switch between cell types in general corresponds to the predicted direction of gene expression change: 6 genes out of 10 in this region have higher expression in SJCRH30 and A375 than in GM12878 and endothelial cells (Fig. 3f).

### Specific sets of variable chromosome interactions correspond to coordinated variations in chromatin accessibility and gene expression

Beyond the role of LADs, to further connect chromosome structure fluctuations to gene regulation and chromatin state, we took advantage of DNase-seq and RNA-seq datasets across many cell types available from ENCODE [18]. DNase-seq profiles chromatin accessibility and whether genomic regions are likely to be open to be regulated or transcribed. On the other hand, RNA-seq represents the combination of transcription, RNA stability, and RNA degradation. In our study, we treated signals measured by RNA-seq as an indicator of transcription. We first calculated chromatin accessibility and transcription variation across all cell types at 100kb bin resolution for each chromosome. Overall, transcription variation has a positive correlation with chromatin accessibility variation (Figure S4). We note that regions that are constitutively LADs or non-LADs (ciLAD and cLAD) tend to have low variation in transcription and chromatin accessibility, while regions that are fluctuating (fiLAD and fLAD) show higher transcription and accessibility variation (Fig. 3d). We next compared regions of high accessibility and transcription variability with the key variable 3D interacting regions identified by E-PCA. We first divided genomic bins into two sets based on transcription or accessibility variation, high and low respectively. Next, we quantified what proportion of key interactions belong to the high variation set. We found that more than 50% of key interactions belong to both high-variation sets in positive E-PC1 and negative E-PC2 and high transcription variation set in negative E-PC2 (Fig. 4a). We conducted a permutation test by shuffling high/low variation sets 100 times to assess the statistical significance of this enrichment. This led us to think that modes of positive E-PC1 and negative E-PC2 may be relevant to regulation in chromatin accessibility and transcription.

**Fig. 4.**
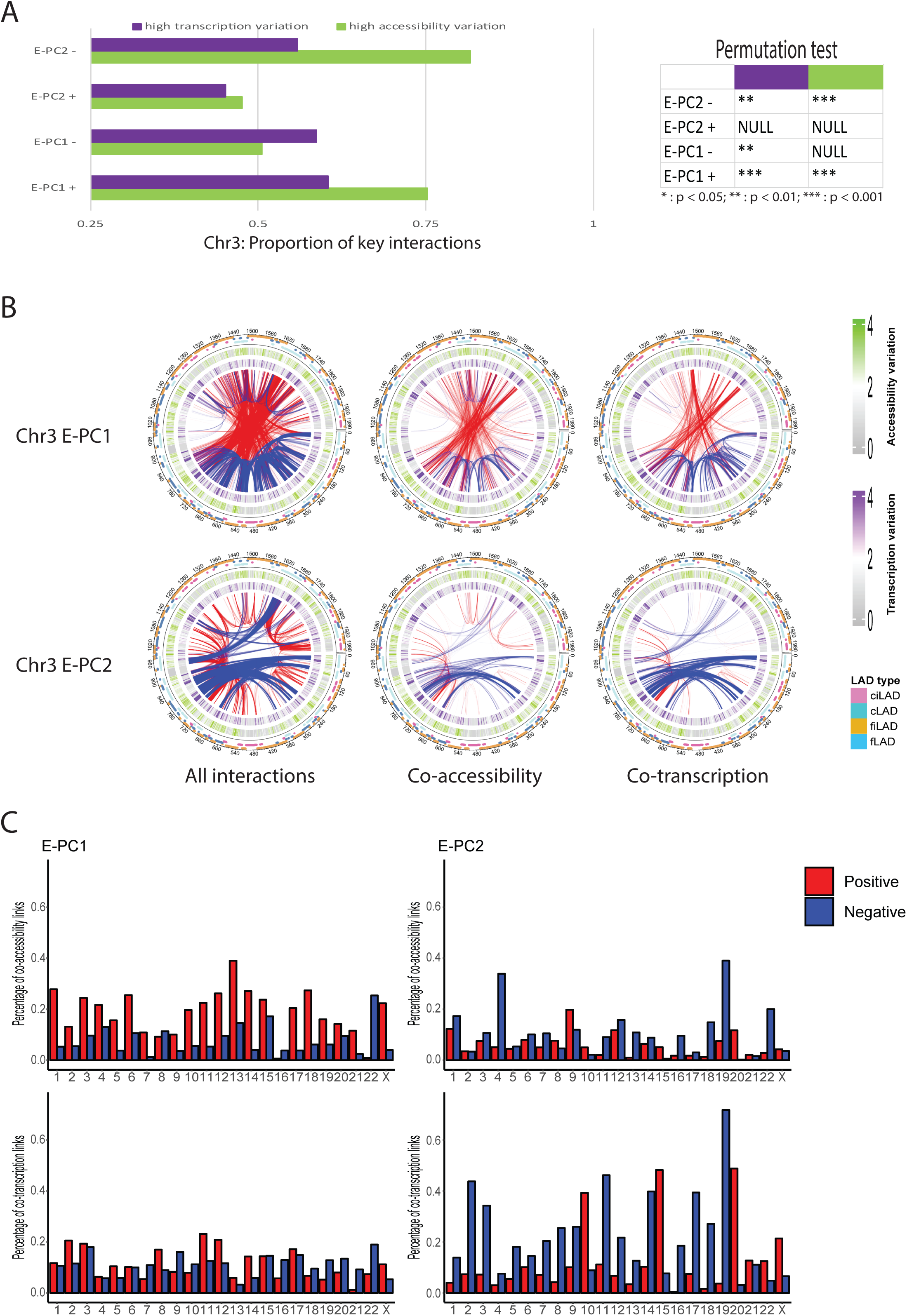
Correlation of interaction sets and chromatin accessibility/transcription variation. a. Percent of key interaction links in high transcription or accessibility variation bins. Right: permutation of high variation bin assignments tests whether key links are statistically enriched in each category. b. Top 1% key variable contacts on chr3 are shown in circular plots for E-PC1 (top row) and E-PC2 (bottom row). LAD type (outer ring), accessibility variation (2^nd^ ring), and transcription variation (3^rd^ ring) are shown around the interaction links. Middle column: interaction links are filtered to keep only those where linked regions show correlated accessibility patterns across cell types. Right column: interaction links filtered to keep only those where genes connected by interactions are co-expressed across cell types. b. Quantification of the % of key interaction links from each PC that are also co-accessible (top row) or co-transcribed (bottom row).

We further hypothesized that pairs of genomic regions connected by key variable 3D interactions would change their transcription and accessibility in a coordinated manner. Indeed, previous work has already shown that correlations in chromatin state between regions can be used to predict the A/B compartment interaction pattern of a single cell types [19]. To examine how coordinated chromatin or transcription relates to key variations in interactions across cell types, we isolated the key interactions that bridge co-transcribed or co-accessible regions, as defined by covariation of bin pairs using RNA-seq and DNase-seq datasets (Methods). We find that certain sets of correlated interactions are highly covariant in accessibility and transcription while other sets are not. For chr3, for example, most of the positive E-PC1 links are also co-variant in accessibility, and most of the negative E-PC2 links are co-variant in transcription (Fig. 4b). Interestingly, we observe that E-PC1 relates to co-accessibility while E-PC2 relates to co-transcription (Fig. 4c). Specifically, the set of positive E-PC1 interactions very often link co-accessible regions, while the set of negative E-PC2 interactions link co-transcribed regions across most chromosomes. For some chromosomes (such as chr19), the same set of interactions (negative E-PC2) link regions that are both co-transcribed and co-accessible. But it is intriguing that in most cases accessibility and transcription variation relate to different modes of genome structure variation. Overall, these results suggest a model in which chromosome folding patterns captured by positive E-PC1 and negative E-PC2 represent interactions among high-variation regions, in terms of chromatin accessibility and transcription. Further, this suggests that the spatial proximity leads to co-transcription and co-accessibility, suggesting a possible regulatory rule that association and dissociation of these interaction sets lead to coordinated changes in transcription or accessibility in these regions.

To further quantify the correlation of interaction variation with transcription and chromatin accessibility variation, we converted 2D interaction links to a 1D profile by quantitating how often each genomic bin was involved in the positive or negative interaction set. The higher the peak is, the more key this genomic bin is to structural variation across the entire chromosome. We then calculated the Pearson correlation between interaction count of each bin and transcription or chromatin accessibility variation. Most regions involved in key E-PCA interactions have a positive correlation with transcription variation and chromatin accessibility variation (Figure S4c). We found co-transcribed/accessible key interactions have an even higher correlation with transcription/accessibility variation than non-co-transcribed/accessible interactions captured by E-PCA (Figure S4c). This suggests that genes which vary the most in accessibility and transcription across cell types are coordinately controlling their expression and chromatin state through forming and breaking spatial contacts between them. Conversely, there is a separate set of interactions that form and break across cell types which do not link co-regulated genes, and we find that these interactions also do not link regions with highly variable gene expression.

### Variable chromosome interaction sets enriched for certain histone modifications

Since we found that a switch between cLAD and ciLAD interactions was a major feature of positive and negative interaction sets for E-PC1 and E-PC2, we hypothesized that these interaction sets would also be characterized by different histone modifications. LADs have generally low gene expression activity and are modified by suppressive markers, like H3K9me2 and H3K9me3. In contrast, inter-LADs are enriched with genes, higher expression activity and active histone marks like H3K4me3 and H3K27ac [15]. To examine how positive and negative interaction sets identified by E-PCA relate to histone modifications, we integrated E-PCA information with ChIP-seq datasets. We aggregated ChIP-seq peaks for H3K4me1, H3K4me3, H3K9me3, H3K9ac, H3K27me3, and H3K27ac across various cell types and matched them with our identified key variable interactions. The result showed that key interactions in positive E-PC1s and negative E-PC2s have positive correlation with H3K9me3 (an inactive mark) and strong negative correlation with the 4 active marks (Fig. 5a). These interaction sets were also found to co-vary in accessibility (above), so the addition or removal of H3K9me3 and the resulting association and dissociation from the lamina might underlie the forming and breaking of these interactions and their accessibility. In contrast, key interactions in negative E-PC1 and positive E-PC2 do not show any distinguishing strong correlations with different histone marks. This is in line with the fact that these same interaction sets are also not strongly associated with co-accessibility or co-transcription.

**Fig. 5.**
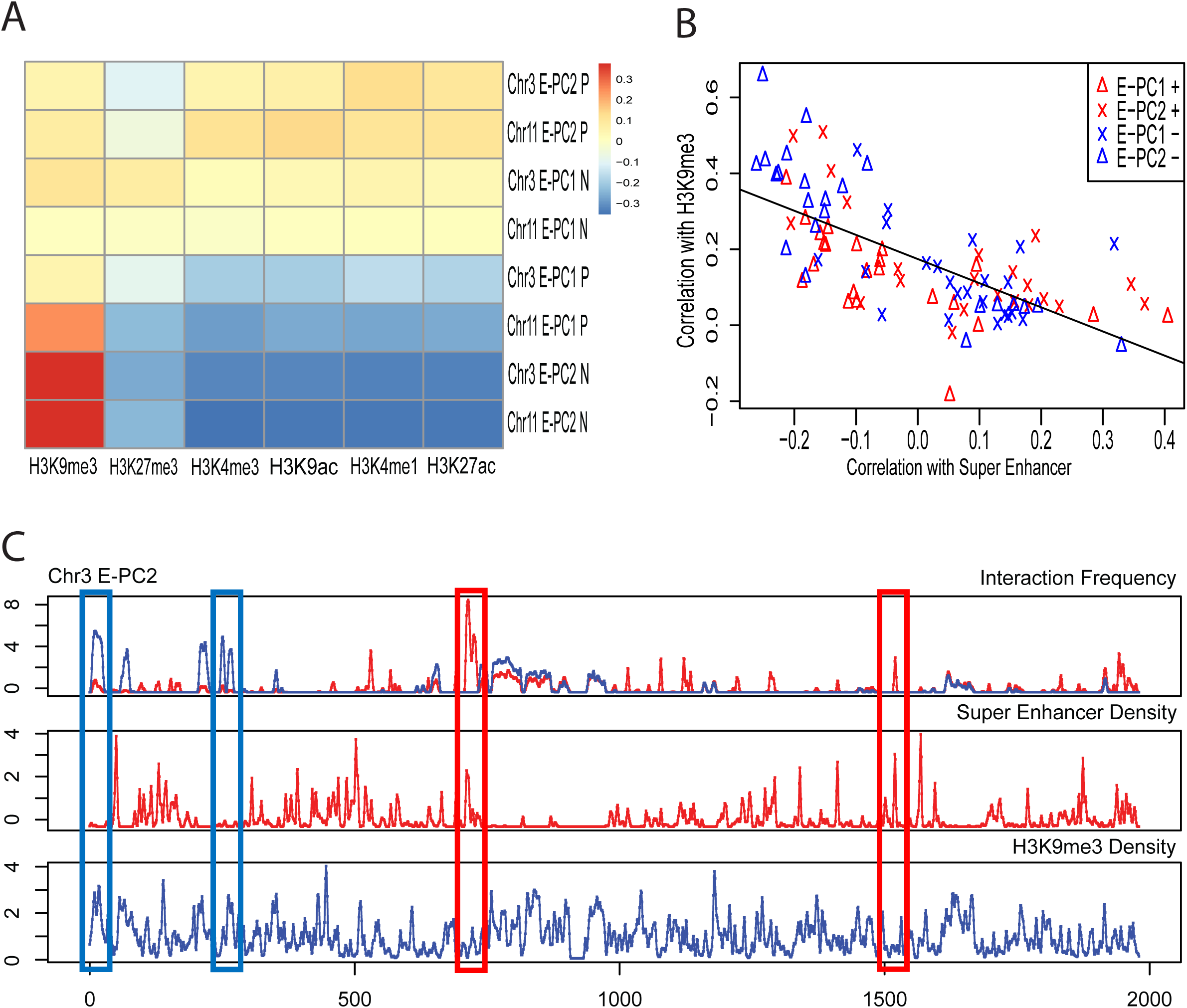
Correlation of interaction sets and histone marks. a. Pearson correlation of participation of each genomic bin in positive or negative interaction sets identified by E-PC1 and E-PC2 of chr3 and chr11 with signal aggregated across cell types for 6 histone marks (H3K4me1, H3K4me3, H3K9me3, H3K9ac, H3K27me3, and H3K27ac). b. Correlation of positive or negative interaction sets with H3K9me3 (Y axis) and super-enhancer (X axis). 92 samples in total from the E-PC1 and E-PC2 across 23 chromosomes. Blue line marks the anti-correlation (Pearson covariance), and grey regions show confidence interval of the fitting line. c. Density peaks of positive (red) and negative (blue) interactions of chr3 E-PC2 (top), super-enhancer (middle), and H3K9me3 (bottom).

Previous studies have shown that super enhancers, which are clusters of enhancers occupied by regulators and mediators, and often defined by high levels of H3K27ac, play a crucial role in shaping cell identity [20], [21]. We therefore evaluated the relationship between our identified key cell type variable Hi-C interactions and super-enhancers. Specifically, we took advantage of previously identified super enhancers by Khan et al. [22]. Their investigation discovered 82,234 super-enhancers in 102 human tissue/cell types. By comparing aggregated super-enhancer peaks with key interactions, we found that key interactions in negative E-PC1s and positive E-PC2s are strongly correlated with super-enhancer peaks, while positive E-PC1 and negative E-PC2 are not. Notably, these sets associated with super-enhancers are opposite from those associated with co-transcription and co-accessibility, described above. Therefore, it appears that spatial clustering of super-enhancer domains forms in opposition to spatial clustering of co-transcription/co-accessible regions (when one set of interactions breaks, the other forms). Accordingly, peaks in negative E-PC1s and positive E-PC2s are strongly correlated with H3K27ac peaks while peaks in positive E-PC1s and negative E-PC2s are strongly correlated with H3K9me3 peaks (Fig. 5b and 5c). Thus, the distinct interaction patterns between positive and negative E-PC values are partly explained by distinctions in histone modification.

### Single cell Hi-C shows that similar principles govern chromosome structure variation both between individual cells and across cell types

Finally, we investigated whether the variations of interaction sets identified across cell types in population averaged Hi-C are related to fluctuations between chromosome structures among single cells. Therefore, we analyzed Hi-C measurements of 6,135 single cells generated by Ramani et al [23]. Due to limited sequencing depth in each individual cell, we considered the interaction sum of every 50 cells as a meta cell. To decide which cells to group together for this sum, we first used Monocle to arrange cells in a pseudotime order [24]. Though this method was first developed to analyze single cell RNA-seq, it was later chosen to order cells by single cell chromatin accessibility [25]. Our results suggest that the algorithm Monocle utilizes is capable of ordering cells by other types of sequencing data, including Hi-C. We first performed E-PCA across meta cells genome wide at 50 Mb resolution. We projected these 6135 single cells to reduced 2-dimensions by Monocle and ordered cells according to pseudotime (Fig. 6a). Next, we grouped every 50-cells as a mega cell in a continuous manner. We reason that grouping cells by the continuous pseudotime will capture different “cell states” that can be represented by the mega cells. Grouping also augments the interaction contacts and we can use E-PCA at a finer resolution. Such a procedure is useful for dealing with single cell Hi-C data that is often “noisy” due sequencing depth issues. Aggregating data across small groups within the dataset combats such noise while still allowing the detection of fluctuation. This idea is guided by the Central Limit Theorem, which describes how the variance is changed with the size of the sub-grouping. Instead of the 100 kb bins we used for bulk Hi-C, we selected 1MB bins for single cell Hi-C E-PCA analysis of the meta cells, and correspondingly used this bin size for LAD annotation and ChIP-seq dataset comparisons.

**Fig. 6.**
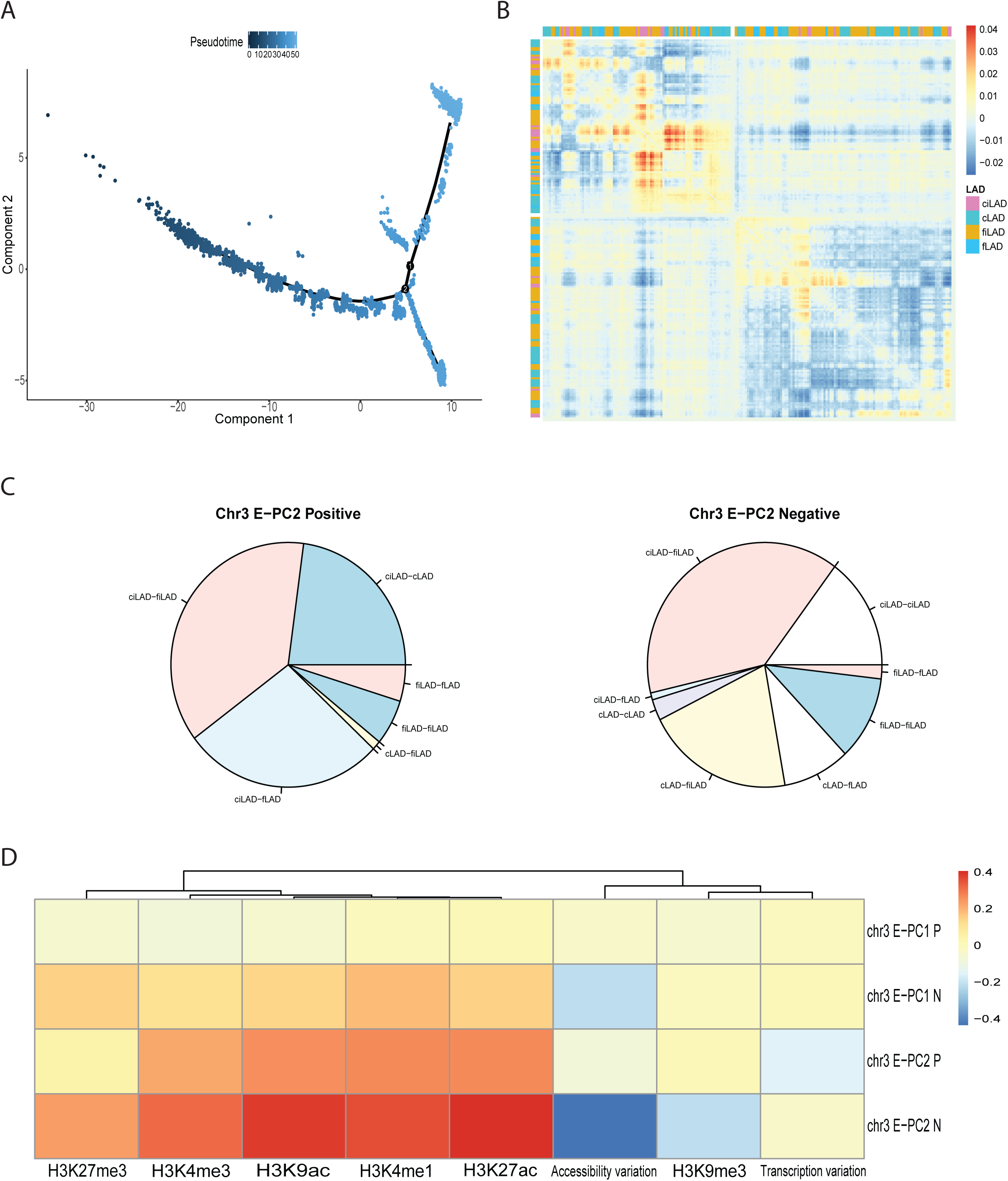
Single cell chromosome structural variation. a. Single cells ordered in 2-dimensional space by Monocle. Cells are ordered in curves and classified into 5 categories by Monocle. b. Major sources of variation across meta-cell Hi-C data from Chr3 identified by E-PC1. c. Pie charts quantify what proportion of each lamina interaction type are found in positive and negative interaction sets, respectively. d. Pearson correlation of single cell Hi-C chr3 positive and negative PC1 and 2 interaction sets with 6 histone marks, accessibility variation and transcription variation.

What we have seen previously hints that the fluctuation of chromosomes between individual cells as observed by imaging showed similar features to the fluctuation of chromosome structures between cell types measured in bulk Hi-C. Here, we found that the first 3 E-PCs from single cell Hi-C do not completely resemble the first 3 E-PCs from bulk Hi-C (Fig. 6b). However, we did see a similarity between bulk Hi-C fluctuations and single cell fluctuations after we annotated interactions with LAD types. We found that the strongest positive and negative interactions in E-PCs captured distinct interaction types associated with LADs. For instance, Chr3 positive E-PC2 interactions tends to involve ciLAD-fLAD interactions. On another hand, Chr3 negative E-PC2 interactions involve more cLAD-fiLAD interactions (Fig. 6c). As in bulk Hi-C, this suggests that a major fluctuation mode between chromosomes in single cells involves lamina association switches. We further found that positive and negative interaction sets in each principal component have distinct correlations with histone marks as well as transcription variation and accessibility variation (Figure S6). Stronger correlation with H3K27ac comes with stronger anti-correlation with H3K9me3, consistent with findings in bulk Hi-C (Fig. 6d).

## Discussion

Our application of E-PCA isolates key sets of chromosome interactions that vary across Hi-C experiments in many cell types. The interaction variations that we are considering are primarily at the A/B compartment scale, both because of our choice of bin size and because we are comparing Hi-C maps in which we have removed the inherent distance decay of interactions with linear genomic distance. The major modes of chromosome contact variation we uncover segregate cell types according to overall lineage and separate cancer from non-cancer cells. This ability to compare numerous Hi-C datasets would allow us in the future to project a new dataset onto an atlas expanded by the E-PC axes. These axes aredefined by an existing range of cell types and the projection would determine whether a perturbation has shifted the genome contacts along one of the key directions identified by the E-PC space. This is conceptually analogous to previously published tracking of cell differentiation or epithelial to mesenchymal transitions along gene expression PCA spaces [26], [27], but would give a picture of whether genome contact changes in such conditions match the expected progression toward chromosome structure characteristics of other cell types.

Beyond being a convenient method to assort cell types along an axis, the interactions revealed as varying in concert across cell types provide valuable conceptual information about what types of chromosome conformation changes are dominant across cell types. Remarkably, the first and second principal components of structure variation are conceptually quite consistent across chromosomes, indicating that they reveal general principles of cell-type specific genome folding. Both modes of variation are dominated by interactions of genomic regions with different characteristic lamin associations. The mode which explains the largest fraction of variation, PC1, involves certain regions changing their interaction partners from repressive LADs to more active interLADs. This suggests that a major mechanism controlling cell identity is sequestering different regions to the nuclear lamina. This result is in line with the recent observations from SPRITE that anchoring to nuclear bodies such as active speckles or the lamina is a major organizing factor of genome structure [28]. Here, we further extend this concept by finding that this is not only true within a single cell type, but that switches in this anchoring are the definitive difference in chromosome structure between cell types. These results also have the interesting implication that there are certain facultative LADs that most often switch their spatial arrangement between cell types. Rather than inventing an entirely new spatial arrangement for each cell type, there appear to be certain regions that are altered within a common framework.

These consistent components of structure variation also represent different aspects of the connection between genome structure and gene regulation. The interactions that form and break in concert captured by PC1 involve regions that also coordinately change their local chromatin accessibility across cell types. Meanwhile, the PC2 interaction set brings together genes that are coordinated in their transcription. Both of these interaction sets show polarized relationships with active and inactive histone modifications, suggesting that histone modifications are involved in the formation of both interaction sets. Often, open chromatin is thought of as synonymous with gene activation. But detailed analyses often find that while the two are correlated, the connection is not one to one. In fact, when accessibility and transcription are measured in the same single cells, the correlations between the two are only partial [25]. Our result suggests that while both local chromatin state and gene expression co-vary with changes in 3D genome folding across cell types, certain interactions might be more involved in coordinated accessibility while others coordinate gene expression.

In both major modes of structure variation, one correlated interaction set is co-accessible or co-transcribed while the anti-correlated set of interactions (which forms when the co-transcribed set breaks, for example) is not related to co-varying genes or highly variable genes across cell types. Strikingly, the set of interactions that co-vary in accessibility and transcription are enriched for H3K9me3, while the set that does not involve interactions between co-varying genes is enriched for superenhancers. This may imply that certain sets of genes that define cell types are marked as heterochromatin and brought together at the nuclear lamina for coordinated repression. On the other hand, cell type specific genes controlled by super-enhancer interactions are activated in some cell lineages and not others but are not co-regulated as a set.

The results of this study raise several questions that need further clarification. First, as this work simply observes co-variation, we cannot prove whether the observed co-variation of chromosome structure, transcription, and chromatin accessibility is critical for maintaining cell identity or the establishment of cell identity. In this study, our information about changes across differentiation are limited, because all samples in the Hi-C data considered here are from differentiated cells. Future applications of E-PCA analyses can investigate correlated chromosome contact changes across differentiation and during specific perturbations that can help tease out the causality of the correlations we observe. Also, the coherent changes we observe occur on average across cell populations. To understand the details of this co-variation, future studies will need to employ techniques like combined single cell sequencing. Joint profiling of chromatin accessibility and gene expression has showed the existence of co-variance to some degree by using single cell technology. Combining single cell Hi-C with RNA-seq or ATAC-seq in the same cells would further clarify the establishment and coherence of cell identity at multiple levels.

## Conclusion

Applying E-PCA across a set of population Hi-C experiments not only allows a comparison of how similar and different genome structure is in these different cell types but uncovers key factors in chromosome structure variation across cell types. We find the major variations in genome folding between different human cell types involve regions that vary in their association with the nuclear lamina. By identifying key variable interaction regions and integrating these with gene expression, chromatin accessibility and ChIP-seq data, we find that co-regulated gene expression corresponds to a certain set of correlated interactions while a different set of interactions is more related to coordinated local chromatin accessibility. Interaction patterns captured by E-PCA are either associated with super-enhancer dense areas or with regions that are co-transcribed/accessible. This suggests some regulatory principles: first, that co-variation in gene expression between sets of highly variable genes across cell types is largely governed by lamina association. In contrast, developmental regulation of specific genes involves interactions with super-enhancers, but not clustering with large sets of co-regulated genes. Both regulatory rules can influence cell-to-cell variation. In this study, we also confirmed that key interaction regions on chromosomes have distinctive properties with respect to histone modifications. These regions either have a high density of active marks, like H3K27ac, or repressive marks, like H3K9me3. Finally, we show that single cells exhibit similar patterns of chromosome structure variation between cells of the same type as are observed between cell types in bulk Hi-C data.

## Methods

### Data sources

LAD type labels were kindly provided by Bas van Steensel [17]. Blood cell type data was obtained with permission from the PCHI-C Consortium that generated this data [29] and the European Genome-phenome Archive. All other datasets can be found on ENCODE Project website (https://www.encodeproject.org/) or were generated from our lab. (See Fig. 2b) Processing of Hi-C matrices followed ENCODE computational guidelines, which can be found at (1); RNA-seq datasets were processed into Bigwig files that contain signals of expression. As for DNase-seq and ChIP-seq, we used signal bed files that present accessibility and histone modification as the form of signal. All Dnase-seq and ChIP-seq signals were generated under ENCODE standard pipeline (2, 3).

1. https://www.encodeproject.org/documents/75926e4b-77aa-4959-8ca7-87efcba39d79/@@download/attachment/comp_doc_7july2018_final.pdf
2. https://www.encodeproject.org/documents/03e5b535-dcdf-4d5e-8ffcfa7bef88b34b/@@download/attachment/DNase-seq%20Pipeline%20Overview.pdf
3. https://www.encodeproject.org/documents/6f6351d4-9310-4a3b-a3c2-70ecac47b28b/@@download/attachment/ChIP-seq_Mapping_Pipeline_Overview.pdf

### Normalization

Normalization of Hi-C matrices followed the guidance described in [30]. We normalized RNA-seq, Dnase-seq and ChIP-seq datasets used in this study by chromosome. For instance, peaks called from one genomic bin were divided by the sum of peaks called from the entire chromosome in the sample and multiplied by the mean of sum of peaks called from the same chromosome across all samples. All normalized datasets were log transformed after the addition of a pseudo-count.

### E-PCA

To calculate the major variations in interaction patterns across Hi-C datasets, we applied the E-PCA method as previously described in the previous study [12]. We use the Spearman correlation matrices (where each entry is the correlation of contact patterns between the given pair of bins) rather than contact frequency matrices in order to emphasize the compartmentalization pattern rather than emphasizing only fluctuations in very local interactions, which may dominate the contact frequency matrix. To cluster E-PCs, we first calculated a Euclidean distance matrix based on the first three E-PC scores of 39 samples and then performed hierarchical clustering.

### Variation/covariation of RNA-seq and DNase-seq

Variations in RNA-seq and DNase-seq were calculated per genomic bin across samples. Then, we divided variation into high and low two sets, by setting median as the threshold for each chromosome. For covariation, we calculated the Pearson correlation for every bin pair, and only correlations larger than 0.5 were defined as co-expression and co-accessibility. To confirm the robustness of the variation and covariation, we performed random down sampling to 75% and repeated over 100 times.

### Key interactions and histone modification

Due to the numbers of possible interactions across the whole genome, we only filtered top 1% interactions in positive and negative E-PCs. This top 1% of interactions represent the most co-varying interactions identified by E-PC. We defined key genomic regions by quantifying the occurrence of one genomic bin in those top 1% interactions. To find key regions that are enriched for a histone modification, we used ENCODE narrowPeak information, binned at 100 kb for each cell type. We then fit Poisson distributions to these binned peak tracks from the same chromosome in the same sample and we selected genomic bins with p < 0.001 and assigned them a binary value of 1. Then, we added up the binarized peaks across all samples. We next calculated the Pearson correlation of the key interaction frequency with key region frequency of each individual histone mark to represent the relation of interactions with histone modifications.

### Single cell Hi-C

We used Monocle to order single cells, as described http://cole-trapnell-lab.github.io/monocle-release/docs/. Different from single cell RNA-seq analysis by Monocle, we chose all interactions instead of variable interaction contacts to order cells. This is because the 50MB genome wide interaction matrix containing 2556 interactions has many fewer dimensions than the single cell RNA-seq expression matrix. After ordering, monocle assigned a pseudotime to each individual cell and we group every 50 cells as a mega cell in a continuous manner to preserve “cell state” at that time point. However, this approach didn’t return a dataset that has the same fine resolution as bulk Hi-C. Therefore, we chose a 1 Mb bin size and performed E-PCA for each chromosome. Due to 10-fold less resolution than bulk Hi-C, we also binned the LAD annotation set, RNA-seq, DNase-seq and ChIP-seq at this lower resolution so we could compare single cell Hi-C to these sequencing data.

## Acknowledgements

We thank Mariano Labrador, Jacob Sanders, Rosela Golloshi, Rebeca San Martin, Emily Stow, and Priyojit Das for insightful discussion. We thank Rosela Golloshi and Jacob Sanders for access to pre-publication Hi-C datasets. The PCHI-C Consortium bears no responsibility for the further analysis or interpretation of these data, over and above that published by the Consortium.

## Figure Legends

**Figure S1**.

Chromosome structural variation by E-PCA across all chromosomes (except chr3, which is presented in Fig. 2). Heatmaps are ordered from E-PC1 to E-PC3 and from chr1 to chrX.

**Figure S2**.

Cell types projected onto the first 2 E-PC axes across all chromosomes (except for chr3, presented in Fig. 2). Scatter plots are ordered from chr1 to chrX.

**Figure S3**.

Quantification of lamina interaction type in key variable interactions. chr21 is presented in figure 3. Pie charts are ordered from E-PC1 to E-PC3 and from Chr1 to ChrX.

**Figure S4**.

Chromatin accessibility variation and transcription variation. a. Scatter plot of variation of accessibility and transcription. Each dot is from one 100kb genomic bin. X and Y axis show how each bin varies with respect to transcription and chromatin accessibility across all cell types. b. Pearson correlation of transcription variation and accessibility variation for each chromosome. c. Relationship between accessibility/transcription variation and types of key interaction links. Each point quantifies one set of interaction links for a given chromosome (ie E-PC2 negative links on chr3). Therefore, there are 92 points in total: from positive and negative E-PC1 and E-PC2 sets across 23 chromosomes. The x-axis represents the correlation between accessibility variation and involvement in this set of links across all bins on the chromosome. That is, a high value indicates that bins involved in this interaction set have more variable accessibility across cell types. A low x-value means this interaction set is unrelated to accessibility variation. The y-axis represents correlation between transcription variation and involvement in the same set of links. Middle and right panels filter key interactions according to whether they link regions that are co-accessible or co-transcribed. Co-accessible/transcribed interaction sets are better correlated with variation in these same features across cell types (right panel). Conversely, some interaction sets that are not co-accessible/transcribed have no relationship to gene expression variability (negative correlations, middle panel).

**Figure S5**.

Correlation of interaction sets and histone marks. Positive or negative interaction sets identified by E-PC1 and E-PC2 of all chromosomes (except chr3 and chr11) correlated with 6 histone marks (H3K4me1, H3K4me3, H3K9me3, H3K9ac, H3K27me3, and H3K27ac).

**Figure S6**.

Correlation of interaction sets and histone marks (single cell Hi-C). Positive or negative interaction sets identified by E-PC1 and E-PC2 of all chromosomes (except chr3) correlated with 6 histone marks (H3K4me1, H3K4me3, H3K9me3, H3K9ac, H3K27me3, and H3K27ac).

**Supplementary Table 1**

Variance of E-PCA. Proportion of variance of each of the first five principal components and the cumulative variance are shown in the table.

## References

1. Klein Allon m, Mazutis L, Akartuna I, Tallapragada N, Veres A, Li V, Peshkin L, Weitz David a, Kirschner Marc w: Droplet Barcoding for Single-Cell Transcriptomics Applied to Embryonic Stem Cells. Cell 2015, 161:1187–1201.

2. Buenrostro JD, Wu B, Litzenburger UM, Ruff D, Gonzales ML, Snyder MP, Chang HY, Greenleaf WJ: Single-cell chromatin accessibility reveals principles of regulatory variation. Nature 2015, 523:486.

3. Assaf R, Oren R, Noam S, Ralph AS, Alon G, David AW, Bradley EB: Single-cell ChIP-seq reveals cell subpopulations defined by chromatin state. Nature Biotechnology 2015, 33.

4. Lieberman-Aiden E, van Berkum NL, Williams L, Imakaev M, Ragoczy T, Telling A, Amit I, Lajoie BR, Sabo PJDMO, Sandstrom R, et al: Comprehensive mapping of long-range interactions reveals folding principles of the human genome.(REPORTS)(Author abstract)(Report). Science 2009, 326:289.

5. Anthony DS, Ming H, Bing R: Genome-wide mapping and analysis of chromosome architecture. Nature Reviews Molecular Cell Biology 2016, 17.

6. Barutcu A, Lajoie B, McCord R, Tye C, Hong D, Messier T, Browne G, van Wijnen A, Lian J, Stein J, et al: Chromatin interaction analysis reveals changes in small chromosome and telomere clustering between epithelial and breast cancer cells. Genome Biology 2015, 16.

7. Rao Suhas sP, Huntley Miriam h, Durand Neva c, Stamenova Elena k, Bochkov Ivan d, Robinson James t, Sanborn Adrian l, Machol I, Omer Arina d, Lander Eric s, Aiden Erez l: A 3D Map of the Human Genome at Kilobase Resolution Reveals Principles of Chromatin Looping. Cell 2015, 162:687–688.

8. Schmitt Anthony d, Hu M, Jung I, Xu Z, Qiu Y, Tan Catherine l, Li Y, Lin S, Lin Y, Barr Cathy l, Ren B: A Compendium of Chromatin Contact Maps Reveals Spatially Active Regions in the Human Genome. Cell Reports 2016, 17:2042–2059.

9. Chathoth KT, Zabet NR: Chromatin architecture reorganization during neuronal cell differentiation in genome. Genome research 2019, 29:613.

10. Bonev B, Mendelson Cohen N, Szabo Q, Fritsch L, Papadopoulos GL, Lubling Y, Xu X, Lv X, Hugnot J-P, Tanay A, Cavalli G: Multiscale 3D Genome Rewiring during Mouse Neural Development. Cell 2017, 171:557–572.e524.

11. Sauerwald N, Kingsford C: Quantifying the similarity of topological domains across normal and cancer human cell types. Bioinformatics 2018, 34:i475–i483.

12. Lindsay RJ, Pham B, Shen T, McCord RP: Characterizing the 3D structure and dynamics of chromosomes and proteins in a common contact matrix framework. Nucleic acids research 2018, 46:8143.

13. Reddy KL, Zullo JM, Bertolino E, Singh H: Transcriptional repression mediated by repositioning of genes to the nuclear lamina. Nature 2008, 452:243.

14. Finlan LE, Sproul D, Thomson I, Boyle S, Kerr E, Perry P, Ylstra B, Chubb JR, Bickmore WA: Recruitment to the Nuclear Periphery Can Alter Expression of Genes in Human Cells (Gene Expression and the Nuclear Periphery). PLoS Genetics 2008, 4:e1000039.

15. van Steensel B, Belmont AS: Lamina-Associated Domains: Links with Chromosome Architecture, Heterochromatin, and Gene Repression. Cell 2017, 169:780–791.

16. Lars G, Ludo P, Emilie B, Wouter M, Marius BF, Wendy T, Bert HE, Annelies De K, Lodewyk W, Wouter De L, Bas Van S: Domain organization of human chromosomes revealed by mapping of nuclear lamina interactions. Nature 2008, 453:948.

17. Kind J, Pagie L, De vries Sandra s, Nahidiazar L, Dey Siddharth s, Bienko M, Zhan Y, Lajoie B, De graaf Carolyn a, Amendola M, et al: Genome-wide Maps of Nuclear Lamina Interactions in Single Human Cells. Cell 2015, 163:134–147.

18. Consortium E: An integrated encyclopedia of DNA elements in the human genome. Nature 2012, 489:57.

19. Fortin J-P, Hansen KD: Reconstructing A/B compartments as revealed by Hi-C using long-range correlations in epigenetic data. Genome biology 2015, 16:180.

20. Whyte Warren a, Orlando David a, Hnisz D, Abraham Brian j, Lin Charles y, Kagey Michael h, Rahl Peter b, Lee Tong i, Young Richard a: Master Transcription Factors and Mediator Establish Super-Enhancers at Key Cell Identity Genes. Cell 2013, 153:307–319.

21. Hnisz D, Abraham Brian j, Lee Tong i, Lau A, Saint-André V, Sigova Alla a, Hoke Heather a, Young Richard a: Super-Enhancers in the Control of Cell Identity and Disease. Cell 2013, 155:934–947.

22. Khan A, Zhang X: dbSUPER: a database of super-enhancers in mouse and human genome. Nucleic acids research 2016, 44:D164.

23. Vijay R, Xinxian D, Ruolan Q, Kevin LG, Frank JS, Christine MD, William SN, Zhijun D, Jay S: Massively multiplex single-cell Hi-C. Nature Methods 2017, 14.

24. Cole T, Davide C, Jonna G, Prapti P, Shuqiang L, Michael M, Niall JL, Kenneth JL, Tarjei SM, John LR: The dynamics and regulators of cell fate decisions are revealed by pseudotemporal ordering of single cells. Nature Biotechnology 2014, 32:381.

25. Cao J, Cusanovich DA, Ramani V, Aghamirzaie D, Pliner HA, Hill AJ, Daza RM, McFaline-Figueroa JL, Packer JS, Christiansen L, et al: Joint profiling of chromatin accessibility and gene expression in thousands of single cells. Science (New York, NY) 2018, 361:1380.

26. Zhang J, Chen H, Li R, Taft DA, Yao G, Bai F, Xing J: Spatial clustering and common regulatory elements correlate with coordinated gene expression.(Research Article). PLoS Computational Biology 2019, 15:e1006786.

27. Rauch A, Haakonsson AK, Madsen JGS, Larsen M, Forss I, Madsen MR, Van Hauwaert EL: Osteogenesis depends on commissioning of a network of stem cell transcription factors that act as repressors of adipogenesis.(Report). Nature Genetics 2019, 51:716.

28. Quinodoz SA, Ollikainen N, Tabak B, Palla A, Schmidt JM, Detmar E, Lai MM, Shishkin AA, Bhat P, Takei Y, et al: Higher-Order Inter-chromosomal Hubs Shape 3D Genome Organization in the Nucleus. Cell 2018, 174:744–757.e724.

29. Javierre BM, Burren OS, Wilder SP, Kreuzhuber R, Hill SM, Sewitz S, Cairns J, Wingett SW, Várnai C, Thiecke MJ, et al: Lineage-Specific Genome Architecture Links Enhancers and Non-coding Disease Variants to Target Gene Promoters. Cell 2016, 167:1369–1384.e1319.

30. Servant N, Varoquaux N, Lajoie BR, Viara E, Chen C-J, Vert J-P, Heard E, Dekker J, Barillot E: HiC-Pro: an optimized and flexible pipeline for Hi-C data processing. Genome biology 2015, 16:259.

